# Meta-analysis of tree diversity effects on the abundance, diversity and activity of herbivores’ enemies

**DOI:** 10.1101/2021.07.05.451117

**Authors:** A. Stemmelen, H. Jactel, E.G. Brockerhoff, B. Castagneyrol

## Abstract

- The natural enemies hypothesis predicts that the abundance and diversity of antagonists such as predators and parasitoids of herbivores increases with the diversity of plants, which can lead to more effective top-down control of insect herbivores. However, although the hypothesis has received large support in agricultural systems, fewer studies have been conducted in forest ecosystems and a comprehensive synthesis of previous research is still lacking.
- We conducted a meta-analysis of 65 publications comparing the diversity, abundance or activity of various groups of natural enemies (including birds, bats, spiders and insect parasitoids) in pure vs. mixed forest stands. We tested the effects of forest biome, natural enemy taxon and type of study (managed vs experimental forest).
- We found a significant positive effect of forest tree diversity on natural enemy abundance and diversity but not on their activity. The effect of tree diversity on natural enemies was stronger towards lower latitudes but was not contingent on the natural enemy taxon level.
- Overall, our study contributes toward a better understanding of the “natural enemies hypothesis” in forest systems and provides new insights about the mechanisms involved. Furthermore, we outline potential avenues for strengthening forest resistance to the growing threat of herbivorous insects.

## Introduction

Forest ecosystems offer a wide variety of ecosystem services that benefit the economy and human wellbeing (Brockerhoff et al., 2017; de Groot et al., 2002). However, in the context of global change, forests are facing increasing threats including herbivory by native and invasive pest species (Grünig et al., 2020; Seidl et al., 2014). It is generally accepted that mixed forests are less vulnerable to insect herbivory than single-species forests (Jactel et al., 2021), a phenomenon commonly referred to as ‘associational resistance’ (Barbosa et al., 2009; Jactel et al., 2017; Letourneau et al., 2011) – though the opposite, ‘associational susceptibility’, may also occur (Schuldt et al., 2014; White & Whitham, 2000). Associational resistance is usually explained by two main mechanisms, driven by the presence of heterospecific neighboring trees: (i) a bottom-up effect (*sensu* Hunter & Price, 1992) on focal host accessibility, with insect herbivores being more likely to find and remain on hosts that are growing in dense single-species stands (the resource concentration hypothesis) and (ii) a top-down effect, corresponding to the control of herbivores population by their natural enemies, supposed to be more abundant in more diverse habitats *(the natural enemies hypothesis)*. The “natural enemies hypothesis” was first presented by Root (1973) and postulates that more diverse habitats provide predators and parasitoids with more abundant and predictable resources (*i.e. prey, breeding and nesting sites, refuge and shelter, etc.*) across spatial and temporal scales, increasing natural enemy abundance and diversity and thus leading to more efficient top-down control of herbivorous pest populations. This hypothesis is well-supported in agricultural systems where it has been shown that crops growing in mixture generally support higher level of predation of herbivores and suffer less loss from herbivory than crops growing in monoculture (Russell, 1989; Letourneau, 1987, Bianchi et al., 2006; Wan et al., 2020). On the other hand, the applicability of the natural enemies hypothesis in forests is still a matter of discussion (Staab & Schuldt, 2020), as studies are reporting positive (e.g. Esquivel-Gómez et al., 2017; Novais et al., 2017), neutral (e.g. Klapwijk & Björkman, 2018; Schuldt et al., 2008) or negative (e.g. Nixon & Roland, 2012) effects of tree diversity on insect parasitism or predation.

Inconsistencies in studies having tested the natural enemies hypothesis in forest ecosystems may arise from multiple sources. Firstly, the natural enemies hypothesis, as it was initially presented, proposed that an increase in abundance and diversity of natural enemies would lead to greater top-down control of herbivores populations. However, relationships between abundance, diversity and actual biological control that natural enemies exert on insect herbivores in forests have not been examined thoroughly. Furthermore, studies having assessed the relevance of the NE hypothesis used different proxies for top-down control by natural enemies, which could lead to different results due to methodological issues. For example, Esquivel-Gómez et al. (2017) tested the effect of tree diversity on spider richness and abundance and found that the *richness* of spiders was significantly higher in mixed stands than in pure stands in a forest of south Mexico. On the contrary, Riihimäki et al. (2005) found no effect of tree species mixing on the *abundance* of spiders in a Norwegian forest. Both studies assessed the response of natural enemies to tree diversity, but used different metrics that does not necessarily covary (i.e. richness vs. abundance) which may have led to the difference in the results observed and in the conclusion made on the natural enemies hypothesis. In recent years, studies using direct measures of predation and parasitism while testing for an effect of tree diversity have increased (Abdala-Roberts et al., 2016; Bellone et al., 2020) but they cannot be directly compared to those measuring only predator abundance. Methodological issues can also arise from the type of study used to test the effect of tree diversity on natural enemies. Tree diversity experiments, where the number of tree species per area is manipulated (e.g in TreeDivNet, Paquette et al., 2018), are used increasingly because they allow for better, more controlled tests of the mechanisms behind natural enemy responses to tree diversity. However, these mainly recent experiments often include mixtures of rather low levels of tree diversity and they typically lack the features of mature forest such as deadwood and diverse microhabitats (Staab & Schuldt, 2020). Therefore, it was hypothesized that these different sources of heterogeneity in studies of natural enemies in mixed forests may make it difficult for clear patterns to emerge (Staab & Schuldt, 2020).

Second, experimental tests of the natural enemies hypothesis in specific locations fail to account for the effect of biogeographical gradients such as latitude, which is known to influence the biodiversity of natural enemies and the strength of various biological interactions (Pennings & Silliman, 2005; Roslin et al., 2017). For example it has been shown that ectothermic arthropod predators such as spiders, ants or carabids are more active under warm climate (Becerra, 2015; Björkman et al., 2011).

Third, it has been found that the effect of tree diversity on natural enemies varied taxa. An experiment conducted in Germany on 150 forest plots showed that the species richness of birds and bats was not related to tree diversity, whereas that of forest-dwelling predatory arthropods was (Penone et al., 2019). Finally, the diet breadth of NE has also been mentioned as an important factor associated with the natural enemies hypothesis in its first mention in the literature (Root et al. 1973). Indeed, it is expected that generalist predators and parasitoids will benefit more from tree diversity than specialists, being able to exploit complementary host or prey resources more efficiently. Therefore, discrepancies could exist between studies conducted in different forest biomes where the proportion of generalists vs. specialists differ, or focusing on different natural enemy taxa of different diet specialization.

In order to quantitatively synthesize the literature testing the natural enemies hypothesis in forest and to better characterize the sources of heterogeneity among studies, we conducted a meta-analysis of 65 published studies from 1993 to 2020 that involved more than 200 case studies. We addressed methodological issues arising from the broad definition of the natural enemies hypothesis by testing whether the effect of tree diversity on natural enemies varied with the metrics used *(abundance, richness and activity)* and the type of study *(tree diversity experiment vs mature forest)*. We also evaluated the importance of the type of natural enemy *(birds, bats, arthropod predators vs. parasitoids)* as well as the forest biome *(tropical, temperate or boreal forest)* to account for the importance of ecological features and biogeographical gradients. Specifically, and albeit the potential effect of the metrics used to assess natural enemy responses to tree diversity, we expect that, following the work of Roslin et al. (2017), the strength of trophic interactions would be higher towards the tropics, which would result in stronger effect of tree diversity on natural enemies at lower latitudes. Additionally, and based on the initial prediction of Root et al. (1973), we expect that more generalist natural enemies would benefit more from the diversity of forests than specialist natural enemies.

## Material & Methods

### Data collection

We conducted an extensive literature search focusing on the effect of tree diversity on natural enemies (hereafter referred to as “NE”) of herbivores. We searched the Web of Science and Scopus databases using the following combination of relevant keywords: [forest OR woodland OR “planted forest”] AND [tree NEAR (diversity OR richness OR composition OR gradients) OR (monoculture OR polyculture OR (pure OR mix*) NEAR stand)] AND [enem* OR predat* OR prey OR parasit* OR top-down OR control* OR bird* OR bat* OR chirop* OR spider* OR ant* OR carab* OR hoverfly OR syrph* OR ladybird OR ichneum* OR hymeno* OR dipter* OR “enemies hypothesis”] AND [diversity OR richness OR abundance OR density OR activity]. We limited the search to papers published in English, between 1950 and 2021. We then screened the title, abstract and full text when appropriate of each previously returned article and included them in the meta-analysis if they met the following criteria:

1. Studies focused on the activity (e.g., predation rate, predation attempts on prey models, percentage of parasitism), abundance or species richness or diversity of NE as a response variable (hereafter “NE response to tree diversity”) in pure vs mixed forest stands or along a gradient of tree species richness or diversity.
2. Studies had to report the mean of the response variable, any measure of variability around means (standard deviation, standard error or 95% confident interval) and the sample size used in the study, or a correlation coefficient between one of the response variables and a gradient of tree species richness or diversity. Wherever necessary, we retrieved that information by digitizing the figures using WebPlotDigitizer version 4.2 (Rohatgi, 2019).

Our literature search yielded a total of 4666 candidate primary studies (last accessed in February 2021), of which we discarded 87% (4083 studies) that did not satisfy our inclusion criteria. We then completed our dataset by checking for any relevant articles present in the cited references of each article retained after the first initial search or in recent reviews. A case study corresponded to any type of NE response to tree diversity, such that there could be more than one case study per primary study. We gave each primary study and case study a single ID. We finally gathered a dataset including 245 case studies from 65 primary studies that were published between 1993 and 2020 across 26 scientific journals (Fig. 1).

**Figure 1.**
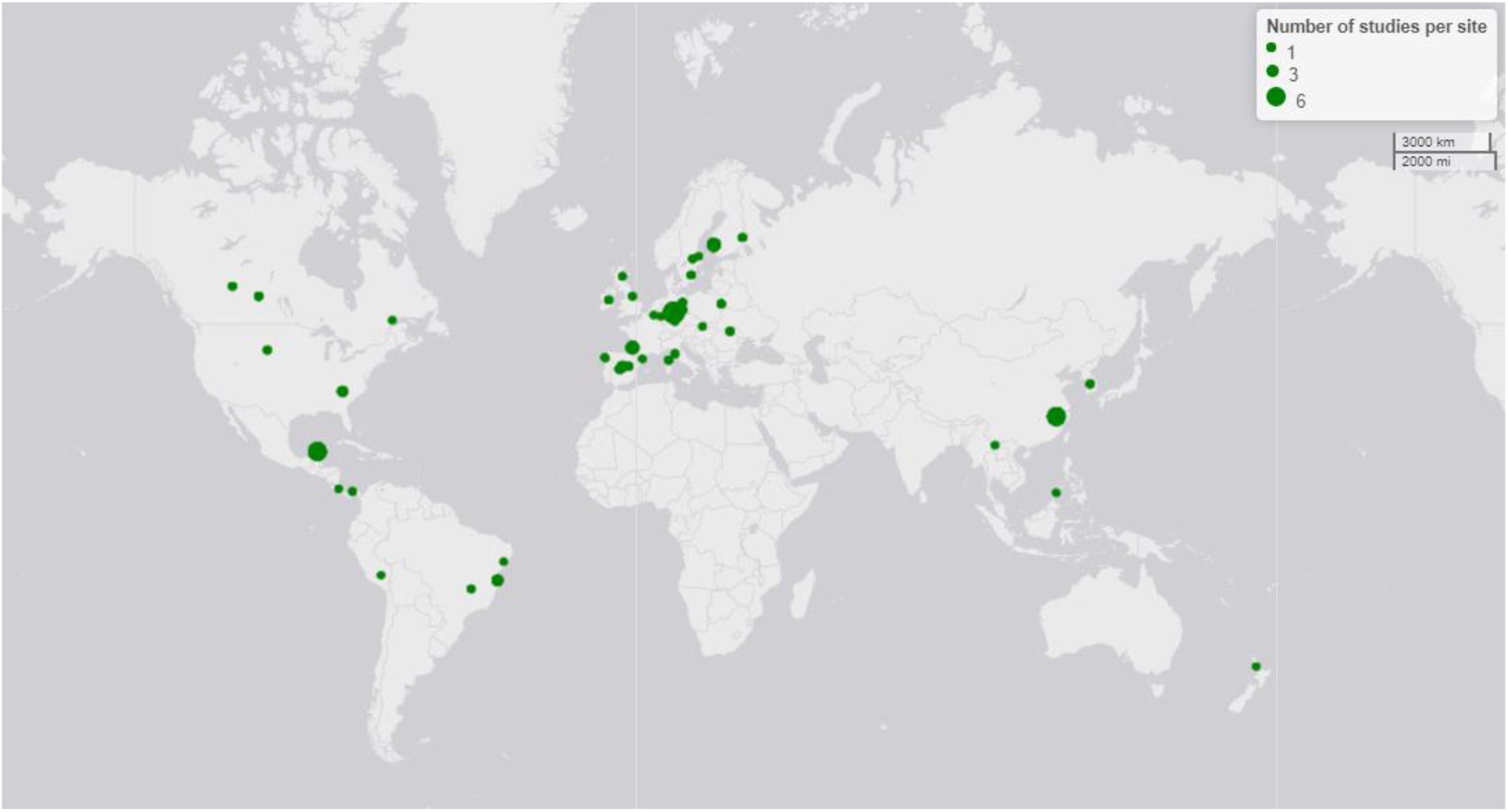
Map of study sites of the 65 articles included in the meta-analysis. Dot sizes are proportional to the number of studies conducted at the same study site.

### Calculating effect sizes

For each case study, we calculated Hedges’ *d* effect size (Hedges & Olkin, 1985) as the standardized differences between NE responses to tree diversity in mixed vs. pure stands. Hedges’ *d* was used because it can account for small sample sizes and uneven variance between control and treatment groups. A positive *d* value corresponds to a positive effect of tree diversity on NE, meaning that NE were more abundant, diverse or active in mixed stands as compared to pure stands. Negative *d* value indicate the opposite. When authors reported correlations between tree diversity and NE responses, we extracted Pearson’s coefficient correlation and sample size from which we calculated Fisher’s z-score. Positive value of Fisher’s z mean that there is a positive correlation between the NE response variable and tree diversity. We then converted Fisher’s *z* to Hedges’ *d* (Koricheva et al., 2013). All effect sizes were calculated with the “metafor” package, using R version 3.6.1 (R Core Team, 2016; Viechtbauer, 2010).

### Moderators

We modeled heterogeneity among effect sizes using four moderators (covariates).

#### Natural enemy response type

We first explored whether the type of response of NE to tree diversity was consistent across several methodological approaches. We categorized NE response as ‘abundance’, ‘diversity’ and ‘activity’. Abundance was assessed using visual identification, trapping or other devices such as ultrasound detectors and included relative and absolute abundance of NE as well as mean number of NE per tree or per trapping device. NE diversity included species richness or Shannon index values. NE activity referred to proportion of real or model hosts or prey that were missing, killed, or had evidence of parasitism or predation attempts.

#### Forest biome

For each primary article, we extracted the geographic coordinates of forest stands, from which we classified each system as ‘boreal’, ‘temperate’ or ‘tropical’ forest ecosystem.

#### Taxa of NE

We distinguished between three categories of NE: Birds and bats (predatory vertebrates), arthropods (generalist predatory invertebrates such as spiders, ants, carabids) and insect parasitoids (specialist predatory invertebrates). When such a classification could not be made from data reported by the authors of primary papers, we categorized NE as ‘unspecified’. When multiple and undiscerned NE taxa were involved, NE were categorized as ‘Multi-taxon’.

#### Type of study

For each study, we classified the forest in which the study was conducted as either “experimental forest”, where the tree diversity gradient was experimentally manipulated (e.g. TreeDivNet), or “natural forest” for other cases.

### Statistical analysis

We first calculated a grand mean effect size (Gurevitch & Hedges, 1999) across all studies to assess the effect of tree diversity on NE. This effect was considered significant if the 95% confidence interval around the grand mean effect size did not include zero. Following Cohen’s guidelines (Cohen, 1988), an effect size of 0.2 or smaller is considered a ‘small’ effect size, 0.5 represents a ‘medium’ effect size and 0.8 or higher a ‘large’ effect size. We estimated consistency among studies by calculating between-study heterogeneity (τ^2^ and associated *Q* statistics) as well as *I^2^*, the standardized estimate of total heterogeneity ranging from 0 to 1 (Koricheva et al., 2013; Nakagawa et al., 2017). As we detected a large amount of residual heterogeneity in the grand mean effect size, we accounted for this using three moderators: NE response type, NE taxa and the forest biome.

We first addressed methodological issues by considering NE response type as a moderator, in interaction with NE taxa or forest biome. The interaction terms were not significant (NE response type × Biome: k = 247, *Q*_M_ = 0.50; P = 0.97; NE response type × NE taxa: k = 247, *Q*_M_ = 3.14, P = 0.53). Additionally, the interaction between NE taxa and forest biome was not significant either (NE taxa × Biome: k = 247, *Q*_M_ = 3.78; P = 0.58). We therefore tested each moderator separately. We reported the results of the omnibus test and interpreted the model coefficients and confidence intervals of each moderator level separately. We considered that model coefficient parameter estimates were significantly different from zero if the 95% CI around each effect size did not overlap zero.

We ran each model using the *rma.mv* function of the “metafor” package and accounted for potential non-independence between case study by including Case ID nested within Study ID as random factors. The use of multiple comparisons to the same control was controlled by using a variance-covariance matrix among effect sizes.

Preliminary analyses testing for an effect of study type (experimental vs natural forest) showed that this factor did not significantly affect NE responses to tree diversity (*Q*_M_ = 0.005, *P* = 0.94) and therefore was not included in the following results section.

Finally, we used several complementary approaches (funnel plot, leave-one out meta-analysis, Rosenberg fail-safe number and cumulative meta-analysis) to evaluate the robustness and sensitivity of our results to several sources of bias (see Appendix A)

## Results

Of the 245 cases studies we analyzed, 159 were associated with arthropods (56 of which were associated with parasitoids), 56 with birds, 16 with bats and 14 with mixed or unidentified NE. NE abundance and richness were assessed 112 and 85 times respectively, while only 48 cases studies assessed NE activity. Studies were mostly conducted in temperate forest (n = 30), followed by tropical (n = 25) and boreal forests (n = 10).

The grand mean effect size (± 95% CI) was significantly positive (0.41 ± [0.17, 0.65], sample size: k = 245, indicating that, overall, NE responded moderately and positively to tree diversity in forest (Fig. 2A. However, there was a large and significant amount of total heterogeneity (τ^2^ = 1.76, Q_T_ = 2899.32, *P* < 0.001), mainly attributed to between-study heterogeneity (I^2^ = 0.91).

**Figure 2.**
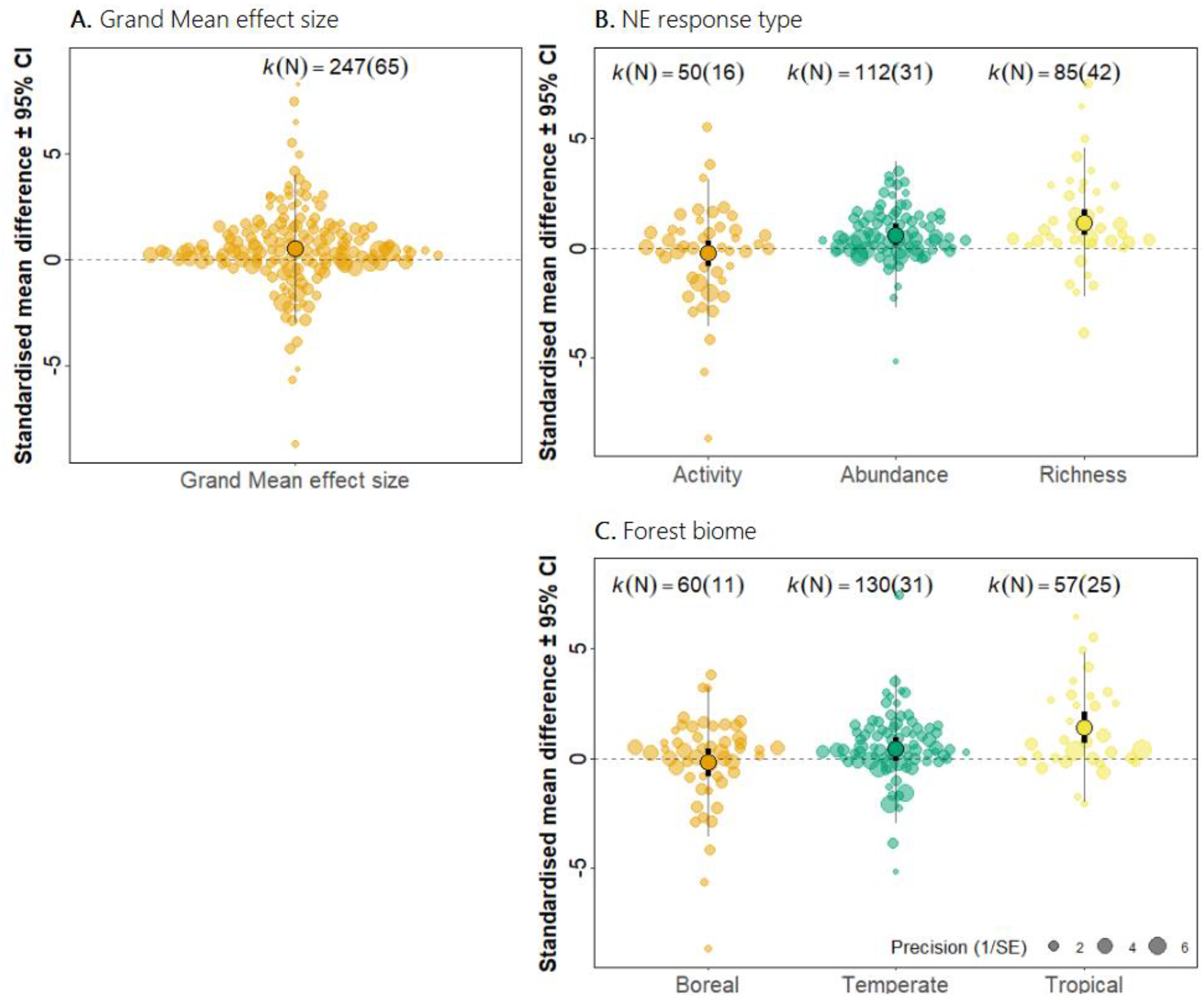
Summary of tree diversity effects on natural enemies (NE). (a) Grand mean effect size. (b) Effect of NE responses type. (c) Effect of the forest biome. Dots and error bars represent model parameter estimates and corresponding 95% confidence intervals. Black circled dots represent the mean effect size for each moderator. Vertical thick lines represent the 95% confidence interval. Thin vertical lines represent the prediction interval, which is the expected range of true effects in similar studies. K (n) represent the number of case studies (K) and the number of primary studies (n) for each moderator level.

We found a significant effect of the metric (i.e. abundance, diversity or activity) used to characterize NE response to tree diversity (*Q*_M_ = 8.34, *P* = 0.015, k = 247). Both NE abundance (0.46 ± [0.14, 0.79]) and diversity (0.68 ± [0.36, 1.01]) were significantly higher in mixed than in pure forests, while NE activity did not significantly influence NE response to tree diversity (−0.16 ± [−0.61, 0.30]) (Fig. 2B).

We found a significant effect of the forest biome in which studies took place (*Q*_M_ = 19.64, *P* = 0.04, k = 247). Mean effect size was large, positive and significantly different from zero in tropical forests (1.43 ± [0.72, 2.14]) but non-significant in temperate forests (0.08 ± [−0.09, 0.95]) and in boreal forests (−0.16 ± [−0.84, 0.50]) (Fig. 2C).

Model coefficient parameter estimates were positive for birds/bats (0.77 ± [0.05, 1.49]) and arthropods (excluding parasitoids) (0.68 ± [0.10, 1.25]), almost null for parasitoids (0.09 ± [−0.78, 0.97]) and negative for “mixed” taxa (−0.56 ± [−1.76, 0.64]). However, the identity of the NE taxon did not influence NE response to tree diversity in forest (*Q*_M_ = 2.11, *P* = 0.54, k = 247).

Finally, we found no significant two or three-way interaction between NE response type, forest biome and NE taxa.

## Discussion

Our meta-analysis provides the first quantitative and up-to-date assessment of the effect of tree species diversity on natural enemies of insect herbivores in forest ecosystems. Overall, NE abundance and diversity were higher in mixed than in pure forests, but not their activity, thus providing positive although partial support to the natural enemies’ hypothesis in forest. This also suggest that increasing forest tree diversity may not automatically lead to stronger top-down control of herbivores by natural enemies and this is corroborated by the large amount of between-study heterogeneity found in our quantitative meta-analysis. Those findings are also consistent with recent studies assessing the effect of tree diversity on natural enemies in forest (reviewed by Staab & Schuldt, 2020) and indicates that this hypothesis may not apply universally in forests. However, we were able to identify some key factors explaining the variation in response of NE to tree diversity in forest.

Consistent with the predictions of the natural enemies hypothesis, we confirm that the abundance and diversity of NE are higher in mixed species forests. Two main non-exclusive mechanisms may explain this pattern. First, diverse forests are expected to host more stable populations of herbivores throughout the year (Lawton & Strong, 1981; Siemann et al., 1998; Vehviläinen et al., 2007) and are thus more likely to provide natural enemies with abundant and various prey and hosts, allowing them to maintain stable populations. In addition, more diverse habitats can offer a better supply of complementary food such as pollen or nectar which can improve the fitness of natural enemies and lead to higher abundance (Cappuccino et al., 1999; Russell, 1989) as well as more microhabitats for nesting or resting (Asbeck et al., 2019; Batáry et al., 2014), and shelter against adverse conditions. In turn, tree diversity had no overall effect on NE activity. One possible ecological explanation could be that intraguild competition, which arises when natural enemies filling the same ecological niche co-occur and share the same prey population (Finke & Denno, 2003; Rosenheim, 1998), or when intraguild predation (e.g. super predators or hyperparasitoids) weakens top-down control of host or prey (Rosenheim et al., 1995), even though the abundance or richness of natural enemies increases in mixed forests. For example, Sanders et al. (2011) found that increasing spider richness did not influence prey control and that the overall outcome strongly depends on the interference between the predators species involved. Another explanation could be that, just as host trees are more difficult to locate and colonize by herbivorous insects in mixed forests because they are physically or chemically hidden by their heterospecific neighbours (Jactel et al. 2021), prey may be more difficult to find by their predators in more complex forests (Nell et al., 2018; Tarbox et al., 2018). Lastly, it is technically much more difficult to properly estimate predation or parasitism of forest insect herbivores than to estimate NE abundance. We could retrieve only 15 papers on NE activity and most of them were based on the use of sentinel caterpillars (38% of all methods used to assess NE activity). There is therefore a need for more studies assessing concomitantly natural enemy abundance/richness and activity in order to properly understand the link between NE abundance and richness and the actual biological control they exert on herbivores.

Natural enemy responses to tree diversity were stronger towards lower latitudes. In a recent review, Staab & Schuldt (2020) already pointed out that latitude and climate could be an important factor shaping NE response to tree diversity in forest and, because tree diversity is much higher in tropical than in temperate and boreal forests, it could have stronger effects on natural enemies. Additionally, previous work highlighted an increase in natural enemy activity or abundance with decreasing latitude (Roslin et al., 2017; Zvereva et al., 2020). Many natural enemies of herbivores are ectothermic arthropods such as spiders, wasps, carabids or ants, and temperature is known to be a key abiotic factors affecting their metabolism and activity (Deutsch et al., 2008; Zvereva & Kozlov, 2006). As such, higher activity of natural enemies, associated with higher overall tree species diversity and stronger trophic interactions for all NE taxa in tropical forests (Gaston, 2000; Hargreaves et al., 2019; Hillebrand, 2004), may explain why the natural enemies responses to tree diversity was stronger in tropical forests than in temperate or boreal forests.

Finally, we found no significant difference of natural enemy responses to tree diversity between NE taxa. Several studies suggest that natural enemy responses to tree diversity are species-specific (Ampoorter et al., 2020; Riihimäki et al., 2005; Staab & Schuldt, 2020; Wan et al., 2020), and some species may not respond to tree diversity, even though the overall effect is positive. For example, Kaitaniemi et al. (2007) found that abundance of predatory ants was higher in mixed forests stands and lead to higher predation of the European pine sawfly, but the abundance of other predators such as spiders and predatory heteropterans did not change between mixed and pure stands. Additionally, the natural enemies hypothesis predicts that generalist natural enemies may benefit more from an increase in tree diversity than specialists, as generalists would be able to make better use of the greater number of alternative prey and host species found in mixed habitats. For example, Legault et al. (2018) found that tree diversity had a positive effect on the generalist parasitoid of the spruce budworm *Apanteles fumiferanae* but did not affect parasitism rate by the specialist parasitoid *Glypta fumiferanae.* Consequently, and given the fact that NE responses to tree diversity can be closely linked to NE identity, it is not surprising that no overall pattern have been found. Finally, it is important to note that studies using multi-taxon approaches to test the responses of different natural enemies to tree diversity under standardized methods were rare (only 6 studies among the 65 included in the meta-analysis). The availability of more such multi-taxon studies would help to unfold global patterns that may otherwise remain unexplored.

## Conclusion and future work

Our results indicate that natural enemy abundance and richness respond positively to an increase in tree diversity in forests, providing at least partial support for the natural enemies hypothesis in forest ecosystems. However, considering that tree diversity does not seem to influence natural enemy activity consistently, it is not yet clear how this effect of tree diversity will affect the ability of natural enemies to exert biological control of herbivore populations. While mixed forests might sustain more abundant and diverse communities of natural enemies, more attention is needed to further understand how this could lead to stronger biotic interactions with their hosts or prey. We therefore recommend that studies assessing NE abundance or richness as a proxy of biological control includes accurate measures of NE activity as well, in order to reach a better understanding of the mechanisms involved in the natural enemies hypothesis in forests.

Finally, we suggest that more attention is given to studies involving multiple taxa or guilds of NE and, ideally that more standardized methods are applied to assess their activity. Such methods could include, among others, the use of dummy caterpillars to estimate predation on lepidopteran-like herbivores or NE exclosure experiments allowing for more precise partitioning of the impact of different NE taxa on herbivorous insect. Otherwise, predicting the role of natural enemies in regulating insect herbivores in diverse forests will remain a challenge.

## Acknowledgments

This study was conducted in the framework of the HOMED project, which received funding from the European Union’s Horizon 2020 research and innovation program under grant 771271. We thank Michael Staab for his friendly review of an early draft of this paper and for his insightful comments.

## Appendix A. Supplementary data

Supplementary data associated with this article can be found, in the online version, at XXXXX. Data are available online from the Data INRAE repository: https://doi.org/10.15454/KS8TLS

