## Supplemental material 1 for "Meta-analysis of tree diversity effects on the abundance, diversity and activity of herbivores’ enemies"

### Supplementary material

#### 1. PRISMA workflow diagram

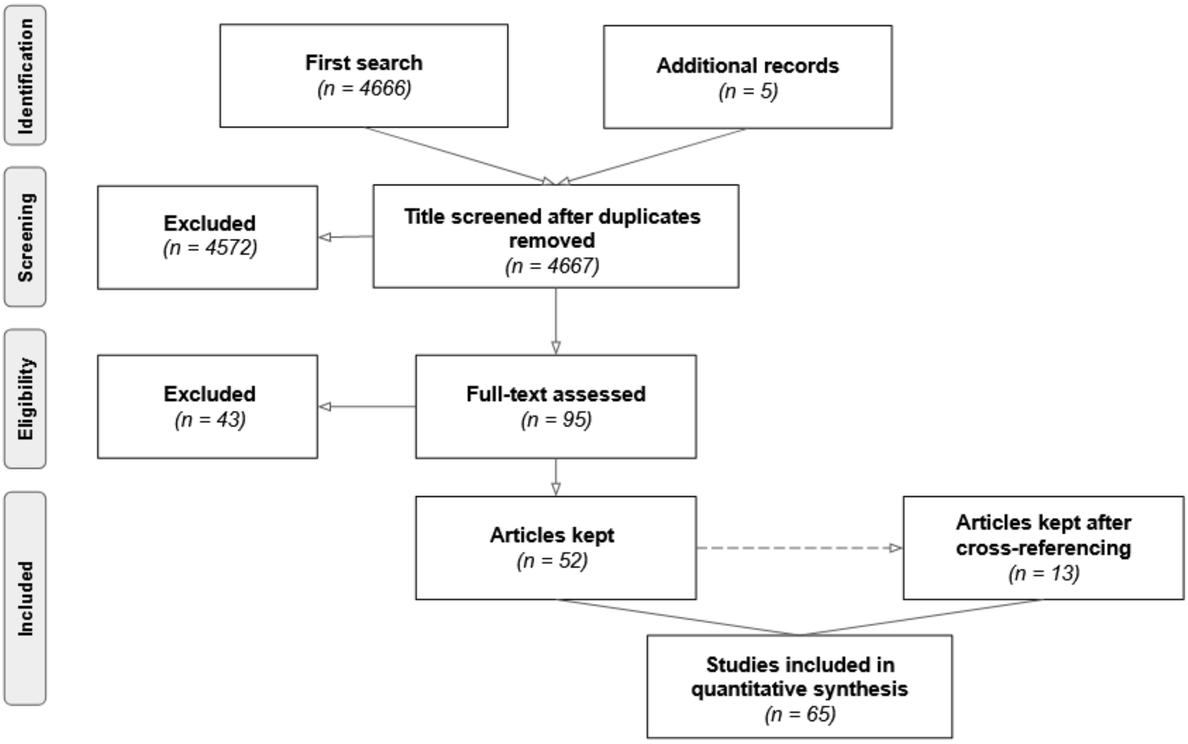


Figure 1. Preferred Reporting Items for Systematic Reviews and Meta-Analyses (PRISMA) workflow diagram

#### 2. Sensitivity analysis and publication bias

We tested the robustness of our results by conducting two complementary sensitivity analyses. First, as some primary studies included more than one study case and could have more influence in the final analysis, we randomly selected one study case per studies and re-ran the analysis with this random dataset. We repeated the process 100 times and checked if the grand mean effects sizes obtained was included in the 95% CI interval calculated on the full dataset (Fig. 2). Second, we checked if the grand mean effect size was particularly influenced by some studies. We recalculated the results of our meta-analysis K-1 times, each times leaving out one study (the "Leave-one-out" method). We calculated a mean effect size for each run and checked that model parameters (mean effect size and 95% CI) were still comparable to the grand mean effect size (Fig. 3). Those analysis indicated that our findings were robust and unbiased.

We addressed publication bias using funnel a plot and the Rosenthal's fail-safe number. The funnel plot created with the 245 study cases was symmetrical, suggesting the absence of publication bias (Fig. 4). Indeed, funnel plots plots effect size estimates (in the x-axis) against a measure of their precision (in the y-axis). It is expected that effect size for less precise studies should scatter more widely on the plot and lead to a funnel shape of the plot. If a bias is present, for example because smaller studies without statistically significant effects are not published, this will lead to an asymmetrical appearance of the funnel plot. In the case of our meta-analyses, studies with lower precision does not tend to show more positive or negative effect size and point are well-distributed around the thick vertical line marking the grand mean effect size. We calculated the fail-safe number as described by Rosenberg (2005). It represents the number of non-significant, unpublished or missing studies that would be needed to make the overall meta-analysis turn non-significant. This number was 1883 and was much greater than the Rosenthal's critical value of 5n+10 (5x63+10=325) (Rosenberg, 2005). Together, those findings suggests no alarming signs of publication bias or abnormalities in the data structure.

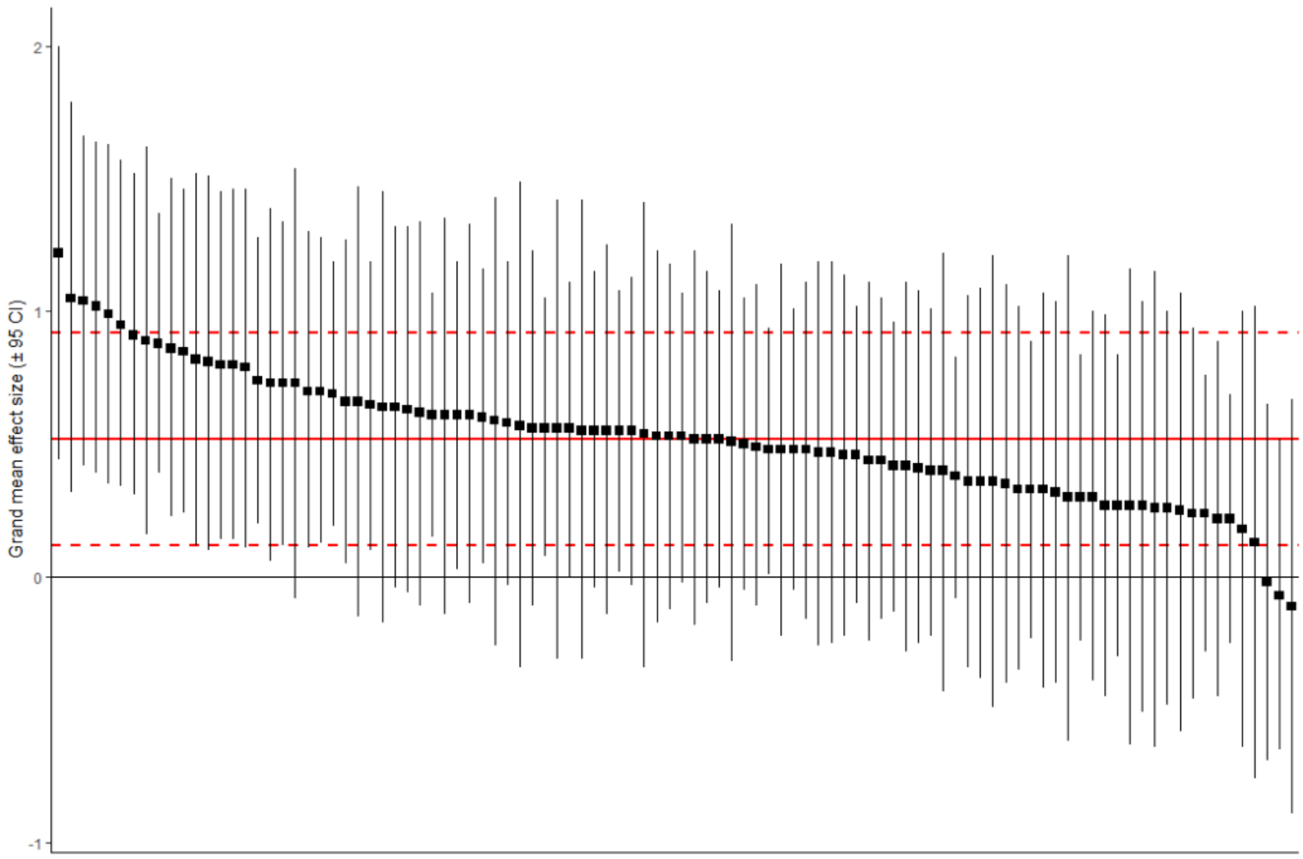


Figure S2. Grand mean effect size ± 95% CI for 100 subset composed of one randomly selected case study per study. Solid and dashed red line represents the Grand mean effect size ± 95% CI calculated on the whole dataset.

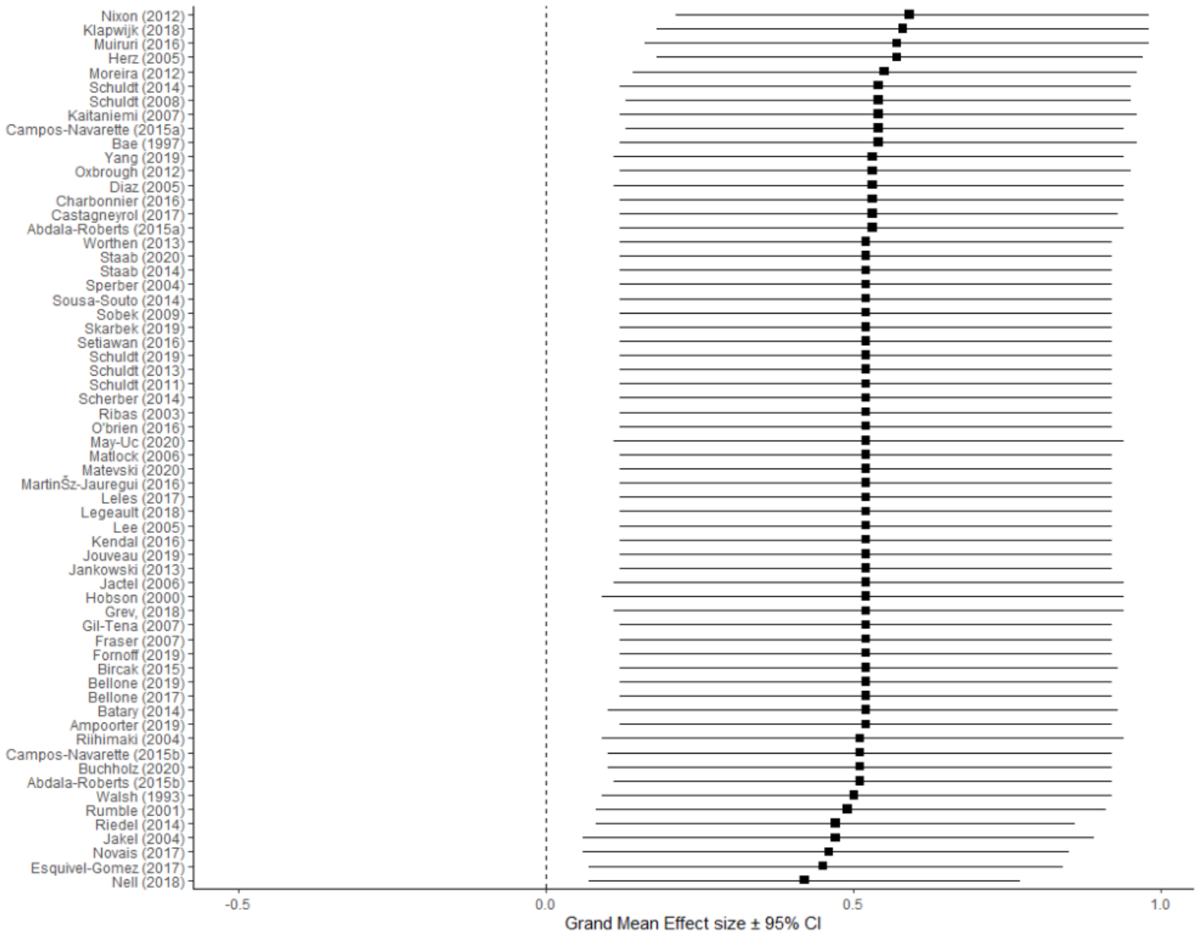


Figure S3. Leave-one-out meta-analysis

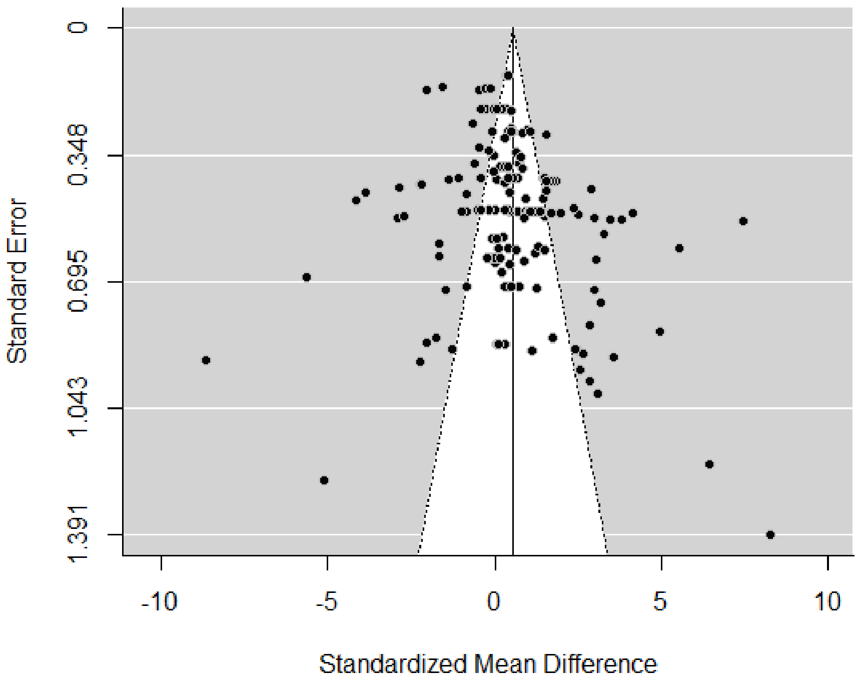


Figure S4. Asymmetric funnel plot representing all 66 studies used in the meta-analysis.

#### 3. List of case studies by moderators

Table 1. Repartition of case studies by moderators

| NE response type | Forest biome | NE taxa |
|---|---|---|
| Activity (n = 50) | Boreal *(n = 32)* | Birds and bats *(n = 12)* |
| | | Parasitoids *(n = 7)* |
| | | Mixed *(n = 13)* |
| | Temperate *(n = 5)* | Birds and bats *(n = 2)* |
| | | Parasitoids *(n = 2)* |
| | | Mixed *(n = 1)* |
| | Tropical *(n = 13)* | Birds and bats *(n = 1)* |
| | | Arthropods *(n = 4)* |
| | | Parasitoids *(n = 7)* |
| | | Mixed *(n = 1)* |
| Abundance (n = 112) | Boreal *(n = 21)* | Birds and bats *(n = 6)* |
| | | Arthropods *(n = 15* |
| | Temperate *(n = 76)* | Birds and bats *(n =17)* |
| | | Arthropods *(n = 28)* |
| | | Parasitoids *(n = 31)* |
| | Tropical *(n = 15)* | Birds and bats *(n = 2)* |
| | | Arthropods *(n = 12)* |
| | | Parasitoids *(n = 1)* |
| Richness (n = 85) | Boreal *(n = 7)* | Birds and bats *(n = 6)* |
| | | Arthropods *(n = 1)* |
| | Temperate *(n = 49)* | Birds and bats *(n = 22)* |
| | | Arthropods *(n = 24)* |
| | | Parasitoids *(n = 3)* |
| | Tropical *(n = 29)* | Birds and bats *(n = 4)* |
| | | Arthropods *(n = 19)* |
| | | Parasitoids *(n = 6)* |

#### 4. List of studies included in the meta-analysis


Abdala-Roberts, L., González-Moreno, A., Mooney, K. A., Moreira, X., González-Hernández, A., & Parra-Tabla, V. (2016). Effects of tree species diversity and genotypic diversity on leafminers and parasitoids in a tropical forest plantation. *Agricultural and Forest Entomology*, *18*(1), 43–51. https://doi.org/10.1111/afe.12132

Abdala-Roberts, L., Mooney, K. A., Quijano-Medina, T., Campos-Navarrete, M. J., González-Moreno, A., & Parra-Tabla, V. (2015). Comparison of tree genotypic diversity and species diversity effects on different guilds of insect herbivores. *Oikos*, *124*(11), 1527–1535. https://doi.org/10.1111/oik.02033

Ampoorter, E., Barbaro, L., Jactel, H., Baeten, L., Boberg, J., Carnol, M., Castagneyrol, B., Charbonnier, Y., Dawud, S. M., Deconchat, M., Smedt, P. D., Wandeler, H. D., Guyot, V., Hättenschwiler, S., Joly, F.-X., Koricheva, J., Milligan, H., Muys, B., Nguyen, D., … Allan, E. (2020). Tree diversity is key for promoting the diversity and abundance of forest-associated taxa in Europe. *Oikos*, *129*(2), 133–146. https://doi.org/10.1111/oik.06290

Bae, W. I., Shin, S. C. (Forestry R. I., & Kim, Z. S. (Korea U. (1997). Difference in occurrence of pine needle gall midge and sucking insects in pure-pine and mixed-pine stands. *FRI Journal of Forest Science (Korea Republic)*. https://agris.fao.org/agris-search/search.do?recordID=KR1998001537

Batáry, P., Fronczek, S., Normann, C., Scherber, C., & Tscharntke, T. (2014). How do edge effect and tree species diversity change bird diversity and avian nest survival in Germany's largest deciduous forest? *Forest Ecology and Management*, *319*, 44–50. https://doi.org/10.1016/j.foreco.2014.02.004

Bellone, D., Björkman, C., & Klapwijk, M. J. (2020). Top-down pressure by generalist and specialist natural enemies in relation to habitat heterogeneity and resource availability. *Basic and Applied Ecology*, *43*, 16–26. https://doi.org/10.1016/j.baae.2019.10.005

Bellone, D., Klapwijk, M. J., & Björkman, C. (2017). Habitat heterogeneity affects predation of European pine sawfly cocoons. *Ecology and Evolution*, *7*(24), 11011–11020. https://doi.org/10.1002/ece3.3632

Birčák, T., & Reif, J. (2015). The effects of tree age and tree species composition on bird species richness in a Central European montane forest. *Biologia*, *70*(11), 1528–1536. https://doi.org/10.1515/biolog-2015-0171

Buchholz, S., Kelm, V., & Ghanem, S. J. (2020). Mono-specific forest plantations are valuable bat habitats: Implications for wind energy development. *European Journal of Wildlife Research*, *67*(1), 1. https://doi.org/10.1007/s10344-020-01440-8

Campos-Navarrete, M. J., Abdala-Roberts, L., Munguía-Rosas, M. A., & Parra-Tabla, V. (2015). Are Tree Species Diversity and Genotypic Diversity Effects on Insect Herbivores Mediated by Ants? *PLOS ONE*, *10*(8), e0132671. https://doi.org/10.1371/journal.pone.0132671

Campos-Navarrete, M. J., Munguía-Rosas, M. A., Abdala-Roberts, L., Quinto, J., & Parra-Tabla, V. (2015). Effects of Tree Genotypic Diversity and Species Diversity on the Arthropod Community Associated with Big-leaf Mahogany. *Biotropica*, *47*(5), 579–587. https://doi.org/10.1111/btp.12250

Castagneyrol, B., Bonal, D., Damien, M., Jactel, H., Meredieu, C., Muiruri, E. W., & Barbaro, L. (2017). Bottom-up and top-down effects of tree species diversity on leaf insect herbivory. *Ecology and Evolution*, *7*(10), 3520–3531. https://doi.org/10.1002/ece3.2950

Charbonnier, Y., Gaüzère, P., van Halder, I., Nezan, J., Barnagaud, J.-Y., Jactel, H., & Barbaro, L. (2016). Deciduous trees increase bat diversity at stand and landscape scales in mosaic pine plantations. *Landscape Ecology*, *31*(2), 291–300. https://doi.org/10.1007/s10980-015-0242-0

Díaz, L. (2006). Influences of forest type and forest structure on bird communities in oak and pine woodlands in Spain. *Forest Ecology and Management*, *223*(1), 54–65. https://doi.org/10.1016/j.foreco.2005.10.061

Esquivel-Gómez, L., Abdala-Roberts, L., Pinkus-Rendón, M., & Parra-Tabla, V. (2017). Effects of tree species diversity on a community of weaver spiders in a tropical forest plantation. *Biotropica*, *49*(1), 63–70. https://doi.org/10.1111/btp.12352

Fornoff, F., Klein, A.-M., Blüthgen, N., & Staab, M. (2019). Tree diversity increases robustness of multi-trophic interactions. *Proceedings of the Royal Society B: Biological Sciences*, *286*(1898), 20182399. https://doi.org/10.1098/rspb.2018.2399

Fraser, S. E. M., Dytham, C., & Mayhew, P. J. (2007). Determinants of parasitoid abundance and diversity in woodland habitats. *Journal of Applied Ecology*, *44*(2), 352–361. https://doi.org/10.1111/j.1365-2664.2006.01266.x

Gil-Tena, A., Saura, S., & Brotons, L. (2007). Effects of forest composition and structure on bird species richness in a Mediterranean context: Implications for forest ecosystem management. *Forest Ecology and Management*, *242*(2), 470–476. https://doi.org/10.1016/j.foreco.2007.01.080

Grevé, M. E., Hager, J., Weisser, W. W., Schall, P., Gossner, M. M., & Feldhaar, H. (2018). Effect of forest management on temperate ant communities. *Ecosphere*, *9*(6), e02303. https://doi.org/10.1002/ecs2.2303

Herz, A., & Heitland, W. (2013). Species diversity and niche separation of cocoon parasitoids in different forest types with endemic populations of their host, the Common Pine Sawfly Diprion pini (Hymenoptera: Diprionidae). *EJE*, *102*(2), 217–224. https://doi.org/10.14411/eje.2005.034

Hobson, K., & Bayne, E. (2000). Breeding Bird Communities in Boreal Forest of Western Canada: Consequences of “Unmixing” the Mixedwoods. *Aspen Bibliography*, *102*. https://doi.org/10.1650/0010-5422(2000)102[0759:BBCIBF]2.0.CO;2

Jactel, H., Menassieu, P., Vetillard, F., Gaulier, A., Samalens, J.-C., & Brockerhoff, E. (2006). Tree species diversity reduces the invasibility of maritime pine stands by the bast scale, Matsucoccus feytaudi (Homoptera: Margarodidae). *Canadian Journal of Forest Research*, *36*, 314–323. https://doi.org/10.1139/x05-251

Jäkel, A., & Roth, M. (2004). Conversion of single-layered Scots pine monocultures into close-to-nature mixed hardwood forests: Effects on parasitoid wasps as pest antagonists. *European Journal of Forest Research*, *123*, 203–212. https://doi.org/10.1007/s10342-004-0030-x

Jankowski, J. E., Merkord, C. L., Rios, W. F., Cabrera, K. G., Revilla, N. S., & Silman, M. R. (2013). The relationship of tropical bird communities to tree species composition and vegetation structure along an Andean elevational gradient. *Journal of Biogeography*, *40*(5), 950–962. https://doi.org/10.1111/jbi.12041

Jouveau, S., Toïgo, M., Giffard, B., Castagneyrol, B., Halder, I. van, Vétillard, F., & Jactel, H. (2020). Carabid activity-density increases with forest vegetation diversity at different spatial scales. *Insect Conservation and Diversity*, *13*(1), 36–46. https://doi.org/10.1111/icad.12372

Kaitaniemi, P., Riihimäki, J., Koricheva, J., & al, et. (2007). *Experimental evidence for associational resistance against the European pine sawfly in mixed tree stands*. *41*, 259–268.

Kendall, L., & Ward, D. (2016). Habitat determinants of the taxonomic and functional diversity of parasitoid wasps. *Biodiversity and Conservation*, *25*. https://doi.org/10.1007/s10531-016-1174-y

Klapwijk, M. J., & Björkman, C. (2018). Mixed forests to mitigate risk of insect outbreaks. *Scandinavian Journal of Forest Research*, *33*(8), 772–780. https://doi.org/10.1080/02827581.2018.1502805

Lee, P.-Y., & Rotenberry, J. (2005). Relationships between bird species and tree species assemblages in forested habitats of eastern North America. *Journal of Biogeography*, *32*, 1139–1150. https://doi.org/10.1111/j.1365-2699.2005.01254.x

Legault, S., & James, P. (2018). Parasitism Rates of Spruce Budworm Larvae: Testing the Enemy Hypothesis Along a Gradient of Forest Diversity Measured at Different Spatial Scales. *Environmental Entomology*, *47*, 1083–1095. https://doi.org/10.1093/ee/nvy113

Leles, B., Xiao, X., Pasion, B. O., Nakamura, A., & Tomlinson, K. W. (2017). Does plant diversity increase top–down control of herbivorous insects in tropical forest? *Oikos*, *126*(8), 1142–1149. https://doi.org/10.1111/oik.03562

Martínez-Jauregui, M., Díaz, M., Sánchez de Ron, D., & Soliño, M. (2016). Plantation or natural recovery? Relative contribution of planted and natural pine forests to the maintenance of regional bird diversity along ecological gradients in Southern Europe. *Forest Ecology and Management*, *376*, 183–192. https://doi.org/10.1016/j.foreco.2016.06.021

Matevski, D., & Schuldt, A. (2021). Tree species richness, tree identity and non-native tree proportion affect arboreal spider diversity, abundance and biomass. *Forest Ecology and Management*, *483*, 118775. https://doi.org/10.1016/j.foreco.2020.118775

Matlock, R., & Edwards, P. (2006). The Influence of Habitat Variables on Bird Communities in Forest Remnants in Costa Rica. *Biodiversity and Conservation*, *15*, 2987–3016. https://doi.org/10.1007/s10531-005-4873-3

May-Uc, Y., Nell, C. S., Parra-Tabla, V., Navarro, J., & Abdala-Roberts, L. (2020). Tree diversity effects through a temporal lens: Implications for the abundance, diversity and stability of foraging birds. *Journal of Animal Ecology*, *89*(8), 1775–1787. https://doi.org/10.1111/1365-2656.13245

Moreira, X., Mooney, K., Zas, R., & Sampedro, L. (2012). Bottom-up effects of host-plant species diversity and top-down effects of ants interactively increase plant performance Proc. *Proceedings. Biological Sciences / The Royal Society*, *279*, 4464–4472. https://doi.org/10.1098/rspb.2012.0893

Muiruri, E. W., Rainio, K., & Koricheva, J. (2016). Do birds see the forest for the trees? Scale-dependent effects of tree diversity on avian predation of artificial larvae. *Oecologia*, *180*(3), 619–630. https://doi.org/10.1007/s00442-015-3391-6

Nell, C. S., Abdala-Roberts, L., Parra-Tabla, V., & Mooney, K. A. (2018). Tropical tree diversity mediates foraging and predatory effects of insectivorous birds. *Proceedings of the Royal Society B: Biological Sciences*, *285*(1890), 20181842. https://doi.org/10.1098/rspb.2018.1842

Nixon, A. E., & Roland, J. (2012). Generalist predation on forest tent caterpillar varies with forest stand composition: An experimental study across multiple life stages. *Ecological Entomology*, *37*(1), 13–23. https://doi.org/10.1111/j.1365-2311.2011.01330.x

Novais, S. M. A., Macedo-Reis, L. E., & Neves, F. S. (2017). Predatory beetles in cacao agroforestry systems in Brazilian Atlantic forest: A test of the natural enemy hypothesis. *Agroforestry Systems*, *91*(1), 201–209. https://doi.org/10.1007/s10457-016-9917-z

O'Brien, M. J., Brezzi, M., Schuldt, A., Zhang, J.-Y., Ma, K., Schmid, B., & Niklaus, P. A. (2017). Tree diversity drives diversity of arthropod herbivores, but successional stage mediates detritivores. *Ecology and Evolution*, *7*(21), 8753–8760. https://doi.org/10.1002/ece3.3411

Oxbrough, A., French, V., Irwin, S., Kelly, T. C., Smiddy, P., & O'Halloran, J. (2012). Can mixed species stands enhance arthropod diversity in plantation forests? *Forest Ecology and Management*, *270*, 11–18. https://doi.org/10.1016/j.foreco.2012.01.006

Pearce, J., Venier, L., McKee, J., Pedlar, J., & Mckenney, D. (2003). Influence of habitat and microhabitat on carabid (Coleoptera: Carabidae) assemblages in four stand types. *Canadian Entomologist - CAN ENTOMOL*, *135*, 337–357. https://doi.org/10.4039/N02-031

Ribas, C. R., Schoereder, J. H., Pic, M., & Soares, S. M. (2003). Tree heterogeneity, resource availability, and larger scale processes regulating arboreal ant species richness. *Austral Ecology*, *28*(3), 305–314. https://doi.org/10.1046/j.1442-9993.2003.01290.x

Riedel, J., Dorn, S., & Mody, K. (2014). Assemblage composition of ants (Hymenoptera: Formicidae) affected by tree diversity and density in native timber tree plantations on former tropical pasture. *Myrmecological News*, *20*, 113–127.

Riihimäki, J., Kaitaniemi, P., Koricheva, J., & Vehviläinen, H. (2005). Testing the enemies hypothesis in forest stands: The important role of tree species composition. *Oecologia*, *142*(1), 90–97. https://doi.org/10.1007/s00442-004-1696-y

Rosenberg, M. S. (2005). The File-Drawer Problem Revisited: A General Weighted Method for Calculating Fail-Safe Numbers in Meta-Analysis. *Evolution*, *59*(2), 464–468. JSTOR.

Rumble, M. A., Flake, L. D., Mills, T. R., & Dykstra, B. L. (2001). Do pine trees in aspen stands increase bird diversity? *In: Shepperd, Wayne D.; Binkley, Dan; Bartos, Dale L.; Stohlgren, Thomas J.; Eskew, Lane G., Comps. Sustaining Aspen in Western Landscapes: Symposium Proceedings; 13-15 June 2000; Grand Junction, CO. Proceedings RMRS-P-18. Fort Collins, CO: U.S. Department of Agriculture, Forest Service, Rocky Mountain Research Station. p. 185-192.*, *18*, 185–192.

Scherber, C., Vockenhuber, E., Stark, A., Meyer, H., & Tscharntke, T. (2014). Effects of tree and herb biodiversity on Diptera, a hyperdiverse insect order. *Oecologia*, *174*. https://doi.org/10.1007/s00442-013-2865-7

Schuldt, A., Both, S., Bruelheide, H., Härdtle, W., Schmid, B., Zhou, H., & Assmann, T. (2011). Predator Diversity and Abundance Provide Little Support for the Enemies Hypothesis in Forests of High Tree Diversity. *PLOS ONE*, *6*(7), e22905. https://doi.org/10.1371/journal.pone.0022905

Schuldt, A., Bruelheide, H., Durka, W., Michalski, S. G., Purschke, O., & Assmann, T. (2014). Tree diversity promotes functional dissimilarity and maintains functional richness despite species loss in predator assemblages. *Oecologia*, *174*(2), 533–543. https://doi.org/10.1007/s00442-013-2790-9

Schuldt, A., Ebeling, A., Kunz, M., Staab, M., Guimarães-Steinicke, C., Bachmann, D., Buchmann, N., Durka, W., Fichtner, A., Fornoff, F., Härdtle, W., Hertzog, L. R., Klein, A.-M., Roscher, C., Schaller, J., von Oheimb, G., Weigelt, A., Weisser, W., Wirth, C., … Eisenhauer, N. (2019). Multiple plant diversity components drive consumer communities across ecosystems. *Nature Communications*, *10*(1), 1460. https://doi.org/10.1038/s41467-019-09448-8

Schuldt, A., Fahrenholz, N., Brauns, M., Migge-Kleian, S., Platner, C., & Schaefer, M. (2008). Communities of ground-living spiders in deciduous forests: Does tree species diversity matter? *Biodiversity and Conservation*, *17*(5), 1267–1284. https://doi.org/10.1007/s10531-008-9330-7

Schuldt, A., & Scherer-Lorenzen, M. (2014). Non-native tree species (Pseudotsuga menziesii) strongly decreases predator biomass and abundance in mixed-species plantations of a tree diversity experiment. *Forest Ecology and Management*, *327*, 10–17. https://doi.org/10.1016/j.foreco.2014.04.036

Setiawan, N. N., Vanhellemont, M., Baeten, L., Gobin, R., De Smedt, P., Proesmans, W., Ampoorter, E., & Verheyen, K. (2016). Does neighbourhood tree diversity affect the crown arthropod community in saplings? *Biodiversity and Conservation*, *25*(1), 169–185. https://doi.org/10.1007/s10531-015-1044-z

Skarbek, C. J., Noack, M., Bruelheide, H., Härdtle, W., Oheimb, G. von, Scholten, T., Seitz, S., & Staab, M. (2020). A tale of scale: Plot but not neighbourhood tree diversity increases leaf litter ant diversity. *Journal of Animal Ecology*, *89*(2), 299–308. https://doi.org/10.1111/1365-2656.13115

Sobek, S., GOßNER, M. M., Scherber, C., Steffan-Dewenter, I., & Tscharntke, T. (2009). Tree diversity drives abundance and spatiotemporal β-diversity of true bugs (Heteroptera). *Ecological Entomology*, *34*(6), 772–782. https://doi.org/10.1111/j.1365-2311.2009.01132.x

Sobek, S., Scherber, C., Steffan-Dewenter, I., & Tscharntke, T. (2009). Sapling herbivory, invertebrate herbivores and predators across a natural tree diversity gradient in Germany's largest connected deciduous forest. *Oecologia*, *160*(2), 279–288. https://doi.org/10.1007/s00442-009-1304-2

Sobek, S., Steffan-Dewenter, I., Scherber, C., & Tscharntke, T. (2009). Spatiotemporal changes of beetle communities across a tree diversity gradient. *Diversity and Distributions*, *15*(4), 660–670. https://doi.org/10.1111/j.1472-4642.2009.00570.x

Sobek, S., Tscharntke, T., Scherber, C., Schiele, S., & Steffan-Dewenter, I. (2009). Canopy vs. understory: Does tree diversity affect bee and wasp communities and their natural enemies across forest strata? *Forest Ecology and Management*, *258*(5), 609–615. https://doi.org/10.1016/j.foreco.2009.04.026

Sousa-Souto, L., Santos, E. D. S., Figueiredo, P. M. F. G., Santos, A. J., & Neves, F. S. (2014). Is there a bottom-up cascade on the assemblages of trees, arboreal insects and spiders in a semiarid Caatinga? *Arthropod-Plant Interactions*, *8*(6), 581–591. https://doi.org/10.1007/s11829-014-9341-0

Sperber, C. F., Nakayama, K., Valverde, M. J., & Neves, F. de S. (2004). Tree species richness and density affect parasitoid diversity in cacao agroforestry. *Basic and Applied Ecology*, *5*(3), 241–251. https://doi.org/10.1016/j.baae.2004.04.001

Staab, M., Liu, X., Assmann, T., Bruelheide, H., Buscot, F., Durka, W., Erfmeier, A., Klein, A.-M., Ma, K., Michalski, S., Wubet, T., Schmid, B., & Schuldt, A. (2021). Tree phylogenetic diversity structures multitrophic communities. *Functional Ecology*, *35*(2), 521–534. https://doi.org/10.1111/1365-2435.13722

Staab, M., Schuldt, A., Assmann, T., & Klein, A.-M. (2014). Tree diversity promotes predator but not omnivore ants in a subtropical Chinese forest. *Ecological Entomology*, *39*(5), 637–647. https://doi.org/10.1111/een.12143

Walsh, P. J., Day, K. R., Leather, S. R., & Smith, A. (1993). The Influence of Soil Type and Pine Species on the Carabid Community of a Plantation Forest with a History of Pine Beauty Moth Infestation. *Forestry: An International Journal of Forest Research*, *66*(2), 135–146. https://doi.org/10.1093/forestry/66.2.135

Worthen, W. B., & Merriman, D. C. G. (2013). Relationships between Carabid Beetle Communities and Forest Stand Parameters: Taxon Congruence or Habitat Association? *Southeastern Naturalist*, *12*(2), 379–386. https://doi.org/10.1656/058.012.0211

Yang, B., Li, B., He, Y., Zhang, L., Bruelheide, H., & Schuldt, A. (2018). Tree diversity has contrasting effects on predation rates by birds and arthropods on three broadleaved, subtropical tree species. *Ecological Research*, *33*(1), 205–212. https://doi.org/10.1007/s11284-017-1531-7
